# Nematicidal and antifeedant activity of ethyl acetate extracts from culture filtrates of *Arabidopsis thaliana* fungal endophytes

**DOI:** 10.1101/2025.01.14.632913

**Authors:** Sandra Díaz-González, María Fe Andrés, Carlos González-Sanz, Soledad Sacristán, Azucena González-Coloma

## Abstract

Endophytic fungi produce a diverse range of bioactive secondary metabolites with potential applications in biopesticide development. This study investigates the nematicidal and antifeedant properties of ethyl acetate extracts from endophytic fungi isolated from wild *Arabidopsis thaliana* populations in Spain. The extracts were tested against the plant-parasitic nematode *Meloidogyne javanica*, and two common insect pests, *Myzus persicae* and *Spodoptera littoralis*. Nine of the 13 extracts demonstrated significant nematicidal and/or antifeedant activity, indicating their potential as biopesticides. The active extracts were derived from six genera: *Alternaria* (3 isolates), *Dydimella* (1), *Dothiora* (1), *Pleiochaeta* (1), *Penicillium* (1), and *Fusarium* (2). Five extracts exhibited nematicidal activity above 90%, with three reducing the total number of *M. javanica* second-stage juveniles hatched from egg masses by 22–37%. Four extracts showed strong settling inhibition (>70%) against *M. persicae*, and three exhibited feeding inhibition against *S. littoralis*. Chemical analysis by GC-MS and LC-MS revealed a wide array of unique secondary metabolites in the active extracts, reflecting substantial chemical diversity, regardless of the fungal origin. This study highlights the potential of fungal endophytes from *A. thaliana* as sources of novel biopesticides, paving the way for future research focused on harnessing these fungi for biopesticide development.

## 1. Introduction

Plant protection products (PPPs) are key agricultural inputs to ensure plant health, crop productivity and food security, since they protect crops from pests and pathogens. However, the inappropriate uses of harmful chemical PPPs can have a negative impact on ecosystems biodiversity and plant, animal and human health. Regulation is nowadays focused on the sustainable and safe production of crops, what has led to a gradual withdrawal of harmful pesticides. In the last decades, alternative approaches to plant protection have been encouraged, such as the use of biopesticides, which tend to pose fewer risks than conventional PPPs^1^. Biopesticides are PPPs derived from naturally occurring compounds or agents obtained from biological sources like animals, plants, and microorganisms^2^. Among microbial sources, fungi play a significant role in biopesticide development due to their production of biologically active secondary metabolites^3,4^. In particular, fungal endophytes - fungi that naturally colonize internal plant tissues without causing disease symptoms^5–7^, are recognized as valuable reservoirs of bioactive compounds, with terpenoids and polyketides being among the most commonly isolated ones. The potential of these metabolites for medicinal and agricultural applications has also been emphasized, highlighting the need for further research in this field^8^. Protection against pests and herbivores is one of the multiple benefits that these microorganisms can naturally provide to the host plant^9^. These microorganisms enhance plant resistance by producing secondary metabolites that deter or inhibit a wide range of pests and herbivores, including microbial pathogens, insects, and nematodes^10,11^. A well-known example is the endophyte *Neotyphodium* sp. which protects plants against herbivores by the production of alkaloids^12^. Given the pressing need for sustainable pest control solutions, increasing attention has been directed toward the bioprospecting of fungal endophytes as potential sources of novel biopesticides^13^.

Plant-parasitic nematodes pose a major threat to global agriculture, causing hundreds of billion annual losses worldwide^14,15^. Among them, root-knot nematodes are particularly destructive, attacking nearly all vascular plants^14^. *Meloidogyne javanica* is one of the most economically important species, serving as a model for studying plant-nematode interactions^16^. Its second-stage juveniles (J2) are the responsible for infecting the plants by penetrating the roots. Infected plants show reduced plant growth and wilting, severely impacting crop productivity^10^. Due to environmental concerns, traditional nematicides such as methyl bromide have been banned or restricted, necessitating the development of sustainable control alternatives^16,17^.

In addition to nematodes, insect pests such as aphids and lepidopterans pose serious threats to agricultural production. Aphids, including *Myzus persicae*, cause significant economic losses through direct feeding and virus transmission, leading to severe yield reductions^18^. *M. persicae* is particularly problematic due to its ability to infest a wide variety of crops and its resistance to multiple insecticide classes, including organophosphates and neonicotinoids^19^. Consequently, biological control methods, including entomopathogens and biological compounds, are being actively explored^19^.

Lepidopteran pests, including *Spodoptera* spp., pose an additional challenge to agriculture^20^. *S. littoralis* is a highly invasive species which damages over 40 plant families, including key crops like wheat, maize, rice, cotton, and vegetables^21^. Its larvae strip leaves and bore into fruits, significantly reducing crop yields^20^. The control of *Spodoptera* spp. requires the massive use of insecticides, since these insects have acquired resistance to all chemical families, including organophosphates, carbamates, and pyrethroids, as well as a more recent family, diamides^22^.This growing resistance further underscores the need for alternative, sustainable pest management strategies.

In this context, the bioprospection of endophytic fungal cultures isolated from different plant species has yielded a wide variety of promising bioactive compounds against nematodes and insect pests^23^. For example, 4-hydroxybenzoic acid, indole-3-acetic acid (IAA) and gibepyrone D produced by the fungal endophyte *Fusarium oxysporum* strain 162 isolated from tomato plants showed high antagonistic effect against the root-knot nematode *Meloidogyne incognita* ^24^. In another study, free fatty acids (oleic, linoleic, palmitic and stearic) present in an extract from the endophytic fungus *Trichoderma* sp. EFI 671, isolated from *Laurus* sp., showed strong antifeedant effects against the aphid *Myzus persicae*^25^. Recent reports on biocidal compounds from fungal endophytes include nematicidal and antifeedant dioxolanones from *Phyllosticta* sp. (YCC4) isolated from *Persea indica* ^26^, aphid antifeedant stempholones from *Stemphylium* sp.^27^ or acaricidal mellein from *Aspergillus*sp.^28^, both isolated from *Bethencourtia palmensis*.

*Arabidopsis thaliana*, a well-established model in plant research, has greatly advanced our understanding of plant–microbe interactions^29,30^. However, studies on its natural fungal endophytes remain scarce. Among the natural associations of this model plant, one notable example is the fungal endophyte *Colletotrichum tofieldiae. C. tofieldiae* establishes a mutualistic relationship with the plant that promotes plant growth and fertility under phosphate-starved conditions^31^. Beyond Arabidopsis, this endophyte also colonizes tomato and maize, improving their growth and yield^32^. Additionally, *C. tofieldiae* has demonstrated biocontrol potential by reducing the prevalence of mycotoxigenic *Aspergillus* spp. and lowering aflatoxin contamination in maize grains^33^. Other studies have identified additional fungal isolates from *Arabidopsis* that provide benefits under stress conditions, including isolates of *Macrophomina*, *Sordaria*, *Phaeosphaeria*, *Chaetomium*, and *Truncatella*^34^. Despite these promising findings, research on the natural fungal endophytes of *A. thaliana* remains limited, particularly regarding their secondary metabolites and potential as bioactive agents against plant pests^30^.

In this study, we explored the biopesticidal potential of fungal endophytes isolated from wild *A. thaliana* plants collected in the central Iberian Peninsula. Culture-filtrate extracts from these endophytes were screened for their nematicidal activity against *M. javanica* and their antifeedant effects on *M. persicae* and *S. littoralis.* The diverse nematicidal and antifeedant effects observed among the extracts emphasize the rich chemical diversity within these endophytes. Our study provides a foundation for exploring endophytes as biotechnological sources of biopesticides, opening new doors for eco-friendly and effective pest management solutions.

## 2. Materials and methods

### 2.1 Fungal strains

The 13 fungal isolates used in this work (Table 1) were isolated in 2010 and 2011 from surface sterilized *A. thaliana* plants from different wild populations of the central Iberian Peninsula described by García et al.^35^.

**Table 1.**
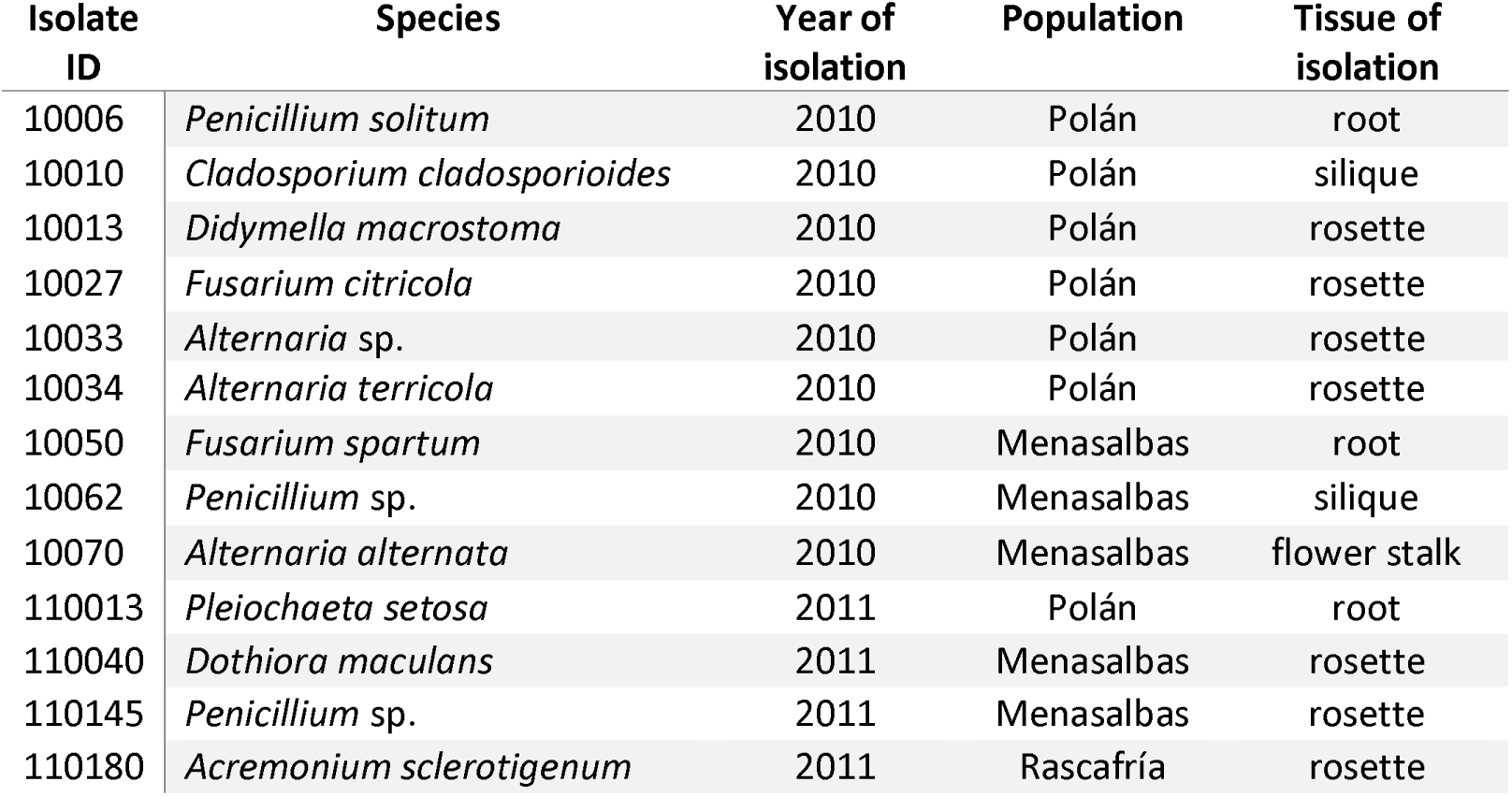
Description of fungal endophytic isolates used for the screening.

| Isolate ID | Species | Year of isolation | Population | Tissue of isolation |
| --- | --- | --- | --- | --- |
| 10006 | *Penicillium solitum* | 2010 | Polán | root |
| 10010 | *Cladosporium cladosporioides* | 2010 | Polán | siliques |
| 10013 | *Didymella macrostoma* | 2010 | Polán | rosette |
| 10027 | *Fusarium citricola* | 2010 | Polán | rosette |
| 10033 | *Alternaria* sp. | 2010 | Polán | rosette |
| 10034 | *Alternaria terricola* | 2010 | Polán | rosette |
| 10050 | *Fusarium spartum* | 2010 | Menasalbas | root |
| 10062 | *Penicillium* sp. | 2010 | Menasalbas | siliques |
| 10070 | *Alternaria alternata* | 2010 | Menasalbas | flower stalk |
| 110013 | *Pleiochaeta setosa* | 2011 | Polán | root |
| 110040 | *Dothiora maculans* | 2011 | Menasalbas | rosette |
| 110145 | *Penicillium* sp. | 2011 | Menasalbas | rosette |
| 110180 | *Acremonium sclerotigenum* | 2011 | Rascafría | rosette |

The identification of each isolate was conducted by sequencing the internal transcribed spacer (ITS) region with the primer pairs ITS1/ITS4^36^. The fungal isolates were restored in potato-dextrose-agar plates (PDA, Difco^TM^) and incubated at room temperature for seven days. Total fungal genomic DNA was extracted from mycelial fragments scraped from fresh fungal culture plates with CTAB method^37^. The PCR conditions were set as follows: 2 min at 94°C for initial denaturation step (1 cycle); 30 s at 94°C, 30 s at 52°C and 1 min at 72°C for amplification cycles (35 cycles); and a final cycle of extension of 10 min at 72°C. Species was assigned based on the best hit of a blast search against Mycobank Database (https://www.mycobank.org/). ITS sequences were aligned with MAFFT v7.525 (Multiple Alignment Fast Fourier Transform)^38^ and trimmed with trimmAl v2.0^39^. Phylogenetic tree was calculated with FastTree v2.1.11^40^ and plotted with iTOL v6^41^.

### 2.2. Culture conditions and liquid-liquid extraction

The fungal isolates were restored in PDA (Difco^TM^) and incubated at room temperature for seven days. Restored colonies were transferred to a new PDA plate and incubated under the same conditions for additional seven days. Spores were scrapped with sterile deionized water from fresh cultures and concentration was determined using a Neubauer Hemocytometer.

The fungal isolates were grown in Malt-Peptone liquid medium (10 g/L Malt Extract, Merck, 2.5 g/L Mycological Peptone, Oxoid^TM^). For each fungal isolate, six 100 mL flasks with a volume of 49 mL of medium each, were inoculated with 1 mL of the fungal spore suspension at the concentration of 5x10^5^ spores/mL, to reach a final concentration of 10^4^ spores/mL of medium. In total, 300 mL of inoculated medium was used per isolate. The flasks were incubated in darkness at 25°C and 120 rpm for 21 days. After incubation, the mycelium was removed from the culture by filtration through a double layer of filter paper with the help of an extraction pump.

Each culture filtrate was subjected to liquid-liquid extraction using ethyl acetate (EtOAc) three times. The volume of the culture filtrate was first measured and the sample was transferred to a separating funnel. An equal volume of EtOAc was added to the funnel, which was then sealed and vigorously shaken to promote mixing. After shaking, the funnel was allowed to stand until phase separation occurred due to the density difference between the aqueous and organic layers. The two phases were then carefully separated by decanting. The EtOAc fraction was subsequently evaporated at 40°C using a rotary evaporator. The dried extract was weighed, and the fraction was stored at 4°C for further use.

### 2.3. Screening

For the initial screening, the extracts were tested in bioassays against *M. javanica* (see section 2.4.1), *M. persicae* (see section 2.5.2), and *S. littoralis* (see section 2.5.3), as described below. Extracts exhibiting mortality rates greater than 90% and inhibiting settling (for *M. persicae*) or feeding (for *S. littoralis*) by more than 70% were selected for further analysis at lower doses. For the *M. javanica* egg hatching assay, the three extracts with the lowest LC50 values were chosen (see section 2.4.2).

### 2.4. Nematicidal activity

#### 2.4.1. Nematicidal Bioassay

The population of *M. javanica* was obtained from Instituto de Ciencias Agrarias, CSIC in Madrid, Spain as described by Moo-Koh et al.^42^. Egg masses of *M. javanica* were handpicked from infected tomato roots. Second-stage juveniles (J2) were obtained from hatched eggs by incubating handpicked egg masses in a water suspension at 25°C for 24 h. The inoculum, was adjusted to a final concentration of 100 J2 nematodes per 95 µL of distilled water. Then, 5 µL of the dissolved extracts or the control solution (DMSO: 0.6% Tween 20) were added to four wells of a 96-well plate containing 95 µL of the nematode suspension, achieving a final extract concentration of 1 µg/µL. Four replicates per treatment were included. The plates were sealed with parafilm to prevent evaporation and were incubated in a growth chamber at 25 ± 1 °C in the dark. Dead J2s were counted at two experimental times (48 and 72 h) using a binocular microscope and mortality rate (M%) was calculated and corrected with Schneider-Orelli’s formula^43^.

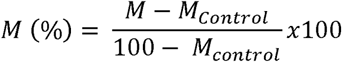

Extracts with a nematicidal activity (M% >90%) were selected for subsequent trials at lower doses (0.5 µg/µL, 0.25 µg/µL and 0.125 µg/µL) to calculate effective lethal doses (LC50 and LC90) at 72 h by probit analysis (software Statgraphics 19, Statgraphics Technologies, Inc.).

#### 2.4.2. Egg hatching inhibition assay

Egg hatching assay was performed as described by Andrés et al.^44^, with the three most active extracts (LC_50_ < 0.5), which were isolates 10034, 10070 and 110040. Four replicates of three egg masses each (a total of 12 egg masses) were tested for each treatment. The three egg masses of each replicate were placed in a well of a 24-well plate and exposed to a total volume of 400 µL of the extracts diluted in DMSO:0.6% Tween 20 at the concentration of 1 µg/µL or DMSO:0.6 % Tween 20 for controls. Plates were sealed with parafilm to prevent evaporation and maintained in a growth chamber in darkness, at 25 ± 1 °C and 70% relative humidity. After five days (day 0) the number of juveniles hatched out of the egg masses was recorded. The test solutions were subsequently removed and the wells with egg masses were washed and filled with sterilized distilled water. Egg hatching was monitored for 4 weeks, until hatching was complete in the control treatment, and then the relative hatching percentages (compared to controls) were recorded.

### 2.5. Antifeedant activity

#### 2.5.1. Maintenance of *M. persicae* and *S. littoralis* colonies

*M. persicae* and *S. littoralis* were reared on bell pepper (*Capsicum annuum* L.) plants and artificial diet^45^, respectively. Host plants together with insect colonies were maintained at 21 ± 2°C, 60-70% relative humidity, and 16 h light: 8 h dark in a growth chamber.

#### 2.5.2. Bioassay of Settling Inhibition of the aphid *M. persicae*

Pepper leaf disks of 2 cm^2^ were cut into two even pieces (1 cm^2^ each). The two leaf sections were set on water-agar (1%) coating the bottom of a ventilated plastic box (3x3x1.5 cm). The dry extracts were redissolved in ethanol at an initial concentration of 10 µg/µL. In each box, one leaf section was treated by spreading 10 µL of the fungal extract over its surface, while the other served as a control, receiving 10 µL of ethanol. Once ethanol evaporated, 10 apterous aphids (24-48 h old) were placed in each plastic box. A total of 20 replicates (boxes) per extract were included in this experiment. The percentage of aphids that settled on each leaf section was recorded after 24 h (at the environmental conditions described above), as described by González-Coloma et al.^46^. Settling inhibition index was calculated at an initial concentration of 100 µg/cm^2^, using the following equation:

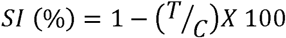

Where:

T = percentage of aphids on the treated section.

C = percentage of aphids on the control section

Extracts with a SI index higher than 70% were considered active and selected for further assays at lower doses (50 µg/cm^2^, 25 µg/cm^2^). EC50 was calculated using a logarithmic regression model with the software Statgraphics 19 (Statgraphics Technologies, Inc.).

#### 2.5.3. Bioassay of Feeding inhibition of *S. littoralis*

Four pepper leaf disks (1 cm^2^) were placed at equal distances on a water-agar (1%) petri dish (9 cm diameter). Two leaf disks were treated with 10 µL of the fungal extract redissolved in ethanol and the other two with 10 µL of the ethanol, as control. After solvent evaporation, two newly molted *S. littoralis* L6 larvae were allowed to feed on the leaf disks at room temperature, until the consumption of either the treated or control disks reached 75%. A total of six replicates (petri dishes) per extract were included in this experiment.

Non-consumed leaf disk area was measured on their digitalized images with the software Image J version 1.53k^47^. Feeding inhibition index was calculated at an initial concentration of 100 µg/cm^2^, using the following equation:

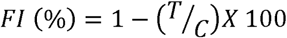

Where:

T = non-consumed areas of treated leaf disks.

C = non-consumed areas of control leaf disks.

Extracts with a FI index higher than 70% were considered active and selected for further assays at lower doses (50 µg/cm^2^, 25 µg/cm^2^). EC_50_ was calculated using a logarithmic regression model with the software Statgraphics 19 (Statgraphics Technologies, Inc.).

### 2.6. Chromatography

#### 2.6.1. Gas Chromatography – Mass Spectrometry

The extracts were dissolved in dichloromethane (DCM) and analyzed by gas chromatography mass spectrometry (GC-MS) using a Shimadzu GC-2010 gas chromatograph coupled to a Shimadzu GCMS-QP2010 Ultra mass detector (electron ionization, 70 eV), equipped with a 30 m × 0.25 mm i.d. capillary column (0.25 μm film thickness) Teknokroma TRB-5 (95%) Dimetil- (5%) diphenylpolisiloxane. Sample injections (1 μl) were carried out by an AOC-20i autosampler. Working conditions were as follows: split ratio (20:1), injector temperature 300°C, temperature of the transfer line connected to the mass spectrometer 250 °C, initial column temperature 110 °C, then heated to 290 °C at 7 °C/min and a Full Scan was used (m/z 35-450). Electron ionization mass spectra and retention data were used to assess the identity of compounds by comparing them with those found in the Wiley 229 and NIST 17 Mass Spectral Database. All extracts (4 μg/μl) were dissolved in 100% DCM for injection.

#### 2.6.2. Liquid Chromatography – Mass Spectrometry

The fungal extracts were analyzed by liquid chromatography coupled to mass spectrometry (LC-MS) in a Shimadzu apparatus equipped with LC-20AD pump and a CTO-10AS VP column oven coupled to a mass spectrometer with triple quadrupole as analyzer (LCMS-8040), with an electrospray ionization source (ESI). An ACE 3 C18 column (150 mm × 4.6 mm, 3 μm particle size) with an ACE3 C18 analytical pre-column was used for the separation. The compounds were eluted with MiliQ water with 0.1% acetic acid (A) and methanol (HPLC-MS grade) with 0.1% acetic acid (B). The solvent gradient started at 38% B reaching 100% in 45 min and 100% during 10 min and then 38% B for 7 min at before the next injection, with a flow rate of 0.5 mL/min. The nitrogen flow (drying gas for solvent evaporation) was 15 L/min. The potential for the electrospray capillary was + 4.50 kV and a Full Scan was used in positive mode (m/z 110– 850) used the Q3 quadrupole with a potential of 1.98 kV and a capillary temperature of 250°C. The heat block temperature was 400°C. The stock solutions of the extracts were injected at 0.5 μg/μl with 5 μl injection through an automatic injector (SIL-20A XR). All extracts (0.5 μg/μl) were dissolved in 100% MeOH for injection.

#### 2.6.3. Hierarchical clustering

Hierarchical clustering of extracts based on GC-MS and LC-MS profiles (% peak area) were calculated based on squared euclidean distance and the complete linkage method with the software Statgraphics 19 (Statgraphics Technologies, Inc.).

## 3. Results

### 3.1. Extraction yields

Different isolates exhibited varying extraction yields of their culture filtrate in ethyl acetate (Supplementary Table S1). Yields (mg of extract per mL of culture filtrate) ranged from 2.6 mg/mL for *A. terricola* (isolate 10034) to 10.4 mg/mL for *D. maculans* (isolate 110040), with an average yield of 5.3 mg/mL across all extracts. Notably, yield differences were observed even among closely related isolates. For instance, *Fusarium* isolates 10027 and 10050 differed by 4.6 mg/mL. *Penicillium* isolates 10006 and 10062 had similar yields (6.9 and 7.8 mg/mL, respectively), whereas isolate 110145 exhibited a lower yield of 2.9 mg/mL (Supplementary Table S1). These findings suggest that extraction yield is isolate-specific rather than genus-dependent.

### 3.2. Nematicidal activity

A total of five extracts out of the 13 tested showed activity (>90% M at 72 h) against *M. javanica* (Table 2 and Table 3). Extract 10034 (*A. terricola*) exhibited the earliest effect causing a mortality rate of 77% at 48 h, while 10033 (*Alternaria* sp.) and 110040 (*D. maculans*) showed the highest effect at 72 h with a 100% of mortality. The most effective extract in terms of lethal dose at 72 h was 10070 (*A. alternata*) with LC_50_ of 0.05 mg and LC_90_ of 0.4 mg. Extract 10034 showed the lowest LC50 at 72 h with 0.04 mg, however LC90 was the second highest.

**Table 2.** Screening for activity of fungal extracts against the plant parasitic nematode *M. javanica* at 1 µg/µL concentration and insect pests *M. persicae* and *S. littoralis* at 100 µg/cm^2^ concentration.

| Isolate/Extract ID | Species | *M. javanica*<br>(%M)<sup>1</sup> | *M. persicae*<br>(%SI)<sup>2</sup> | *S. littoralis*<br>(%FI)<sup>3</sup> |
| --- | --- | --- | --- | --- |
| 10006 | *Penicillium solitum* | 23.2 ± 2.3 | 93.6 ± 2.2 | 72.0 ± 7.9 |
| 10010 | *Cladosporium cladosporioides* | 56.6 ± 3.2 | 59.2 ± 11 | 51.5 ± 10.0 |
| 10013 | *Didymella macrostoma* | 25.1 ± 5.2 | 68.8 ± 5.3 | 81.6 ± 16.4 |
| 10027 | *Fusarium citricola* | 15.5 ± 4.3 | 80.0 ± 7.8 | 31.4 ± 13.3 |
| 10033 | *Alternaria* sp. | 100 ± 0.0 | 63.6 ± 6.2 | 19.8 ± 6.5 |
| 10034 | *Alternaria terricola* | 90.2 ± 2.0 | 52.8 ± 9.6 | 29.6 ± 16.0 |
| 10050 | *Fusarium spartum* | 3.4 ± 2.1 | 70.1 ± 6.4 | 17.3 ± 7.8 |
| 10062 | *Penicillium* sp. | 1.1 ± 0.9 | 57.9 ± 7.7 | 42.2 ± 12.5 |
| 10070 | *Alternaria alternata* | 96.7 ± 0.9 | 80.6 ± 8.1 | 99.4 ± 0.4 |
| 110013 | *Pleiochaeta setosa* | 90.3 ± 3.1 | 64.2 ± 8.5 | 22.6 ± 7.3 |
| 110040 | *Dothiora maculans* | 100 ± 0.0 | 59.8 ± 7.7 | 49.3 ± 15.1 |
| 110145 | *Penicillium* sp. | 39.7 ± 5.8 | 52.3 ± 9.2 | 4.15 ± 2.9 |
| 110180 | *Acremonium sclerotigenum* | 0.0 ± 1.4 | 59.4 ± 10.5 | 17.3 ± 10.6 |

<sup>1</sup>Mortality rate at 72 h. Extracts were considered active when they showed >90% mortality rate at 72 h.
<sup>2</sup>Percent settling inhibition (%SI, n = 200 insects). Extracts were considered active when they showed >70% SI.
<sup>3</sup>Percent feeding inhibition (% FI, n = 12 insects). Extracts were considered active when they showed >70% FI.
Data shows mean ± standard error.

**Table 3.** Mortality rates and lethal doses (LC50 and LC90) of the active ^1^ fungal extracts against *M. javanica in vitro*.

| Extract ID | Species | % Mortality<sup>2</sup> |  | Lethal doses at 72 h (mg) |  | P-value<sup>3</sup> |
| --- | --- | --- | --- | --- | --- | --- |
|  |  | 48 h | 72 h | LC<sub>50</sub> | LC<sub>90</sub> |  |
| 10033 | *Alternaria* sp. | 47.2 ± 3.1 | 100 ± 0.0 | 0.51 | 0.87 | <0.001*** |
| 10034 | *Alternaria terricola* | 77.6 ± 1.8 | 90.2 ± 2.0 | 0.04 | 1.03 | <0.001*** |
| 10070 | *Alternaria alternata* | 3.3 ± 1.1 | 96.7 ± 0.9 | 0.05 | 0.4 | <0.001*** |
| 110013 | *Pleiochaeta setosa* | 27.3 ± 2.9 | 90.3 ± 3.1 | 0.91 | 1.53 | <0.001*** |
| 110040 | *Dothiora maculans* | 56.8 ± 2.7 | 100 ± 0.0 | 0.42 | 0.8 | <0.001*** |

<sup>1</sup>Extracts were considered active when they showed >90% mortality rate at 72 h.
<sup>2</sup>Data shows mean ± standard error.
<sup>3</sup>P values from Probit analysis used for the calculation of LC<sub>50</sub> and LC<sub>90</sub> at 72 h

The active extracts with the lowest LC50 values (10034, 10070, and 110040) were selected to assess their effect on egg hatching. The three treatments resulted in a reduction in the total number of J2 that hatched from egg masses in comparison to controls, being the effect stronger during the first seven days (Figure 1 and Supplementary Table S2), with reductions between 11% and 62% at day 0 (after five days of exposition to extract) and between 41% and 52% at day 7. Extract 10070 showed the highest total hatching inhibition rate with 37%, followed by 110040 with 28% and 10034 with 22%.

**Figure 1.**
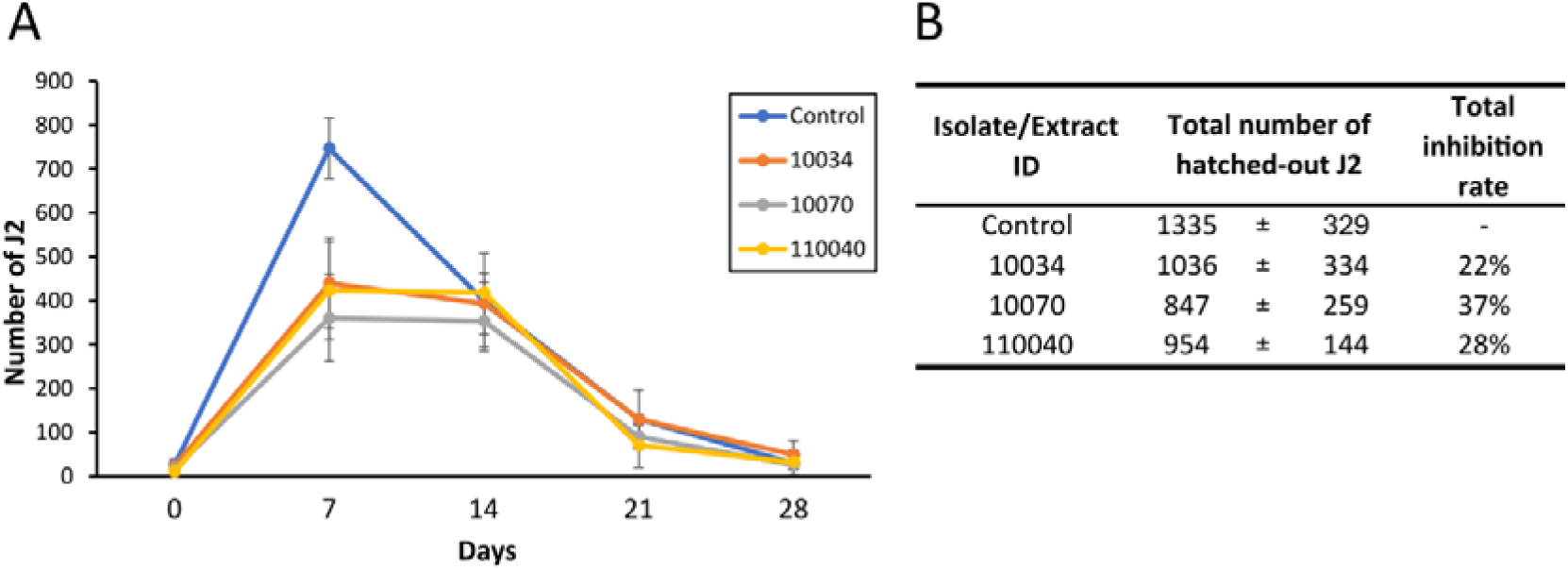
Egg hatching inhibition assay. A) Temporal series of the number of second-stage juveniles (J2) hatched out of three egg masses. B) Total number of J2 hatched out at the end of the experiment and total inhibition rate of the most active extracts, from isolates 10034 (*Alternaria terricola*), 10070 (*Alternaria alternata*) and 110040 (*Dothiora maculans*). Egg masses were exposed for five days to the extracts diluted in DMSO:0.6% Tween 20 or DMSO:0.6 % Tween 20 for controls. The number of J2 hatched out of the egg masses was recorded at day 0 (after five days of exposition to extracts) and at 7, 14, 21 and 28 days immersed in water. Data (n = 4) shows means ± standard deviations.

### 3.3. Antifeedant activity

The extracts from the 13 endophytic fungi were screened for their biocontrol properties against the insect pests *M. persicae and S. littoralis.*Four fungal extracts (10006, 10027, 10050 and 10070), representing 31% of the tested extracts, showed high (>70%) SI effect against *M. persicae* at a starting concentration of 100 µg/cm^2^ (Table 2). With these active extracts we calculated EC_50_ by conducting the assays at lower concentrations (Table 4). Extract from isolate 10050 (*F. spartum*) exhibited the lowest EC_50_ at 20 µg/cm^2^, followed by 10006 (*P. solitum*) with a EC_50_ of 30 µg/cm^2^. The latter was also the extract showing the highest SI rate at 100 µg/cm^2^ with 93.6%.

**Table 4.** Half maximal effective concentration (EC _50_) of fungal extracts with antifeedant activity.

| Extract ID | Species | *M. persicae* |  | *S. littoralis* |  |
| --- | --- | --- | --- | --- | --- |
|  |  | EC<sub>50</sub> (µg/cm<sup>2</sup>) | P value<sup>1</sup> | EC<sub>50</sub> (µg/cm<sup>2</sup>) | P value<sup>1</sup> |
| 10006 | *Penicillium solitum* | 30 | <0.001*** | 76 | 0.105 |
| 10013 | *Didymella macrostoma* | - | - | 60 | 0.001** |
| 10027 | *Fusarium citricola* | 37 | 0.001** | - | - |
| 10050 | *Fusarium spartum* | 20 | 0.007** | - | - |
| 10070 | *Alternaria alternata* | 35 | <0.001*** | 42 | <0.001*** |

<sup>1</sup>P values from exponential regression models used for the calculation of EC<sub>50</sub>.

Three fungal extracts (10006, 10013 and 10070) exhibited high FI effect against *S. littoralis* (Table 2). The extract from isolate 10070 (*A. alternata*) was the most effective with 99.4% FI at 100 µg/cm^2^ and a EC_50_ of 42 µg/cm^2^.

### 3.4. Metabolomic analysis

As a result of the GC-MS analysis, the presence of 117 compounds (>%1 abundance) was revealed, among which only 42 (36%) could be identified (Supplementary Table S3). The number of compounds in each extract ranged from 9 (isolate 10006, *P. solitum*), to 25 (isolate 110013, *P. setosa*), and the presence of the different compounds was mostly specific of each isolate, with just one compound (No. 100: unidentified) present in the extracts of all nine isolates and three compounds (No. 34: 2,10 bisaboladien-1-one (IUPAC: (2E,10E)-bisabola-2,10-dien-1-one); No. 48: 9-Octadecen-1-ol, (Z)- (IUPAC: (Z)-9-octadecen-1-ol), and No. 69: Tributyl acetylcitrate, IUPAC: Tri-n-butyl 2-acetoxypropane-1,2,3-tricarboxylate) shared by the extracts of five or more isolates. Namely, compounds (No. 34) 2,10-bisaboladien-1-one and (No. 69) Tributyl acetylcitrate were present in all three extracts with activity against *M. persicae* (Supplementary Table S3). Samples could be grouped in three clusters according to their volatile-lipid composition (Figure 2A), with cluster 3 divided further in three subgroups. Clusters GC1 and GC2 grouped the extracts from isolates from Polán population (roots and rosette respectively). Cluster GC3 grouped extracts from isolates from the two populations and different plant organs. The grouping did not correlate with the taxonomy of the isolates or the activity (Figure 2C), indicating that the volatile-lipidic composition of the extracts was not specific for these traits.

**Figure 2.**
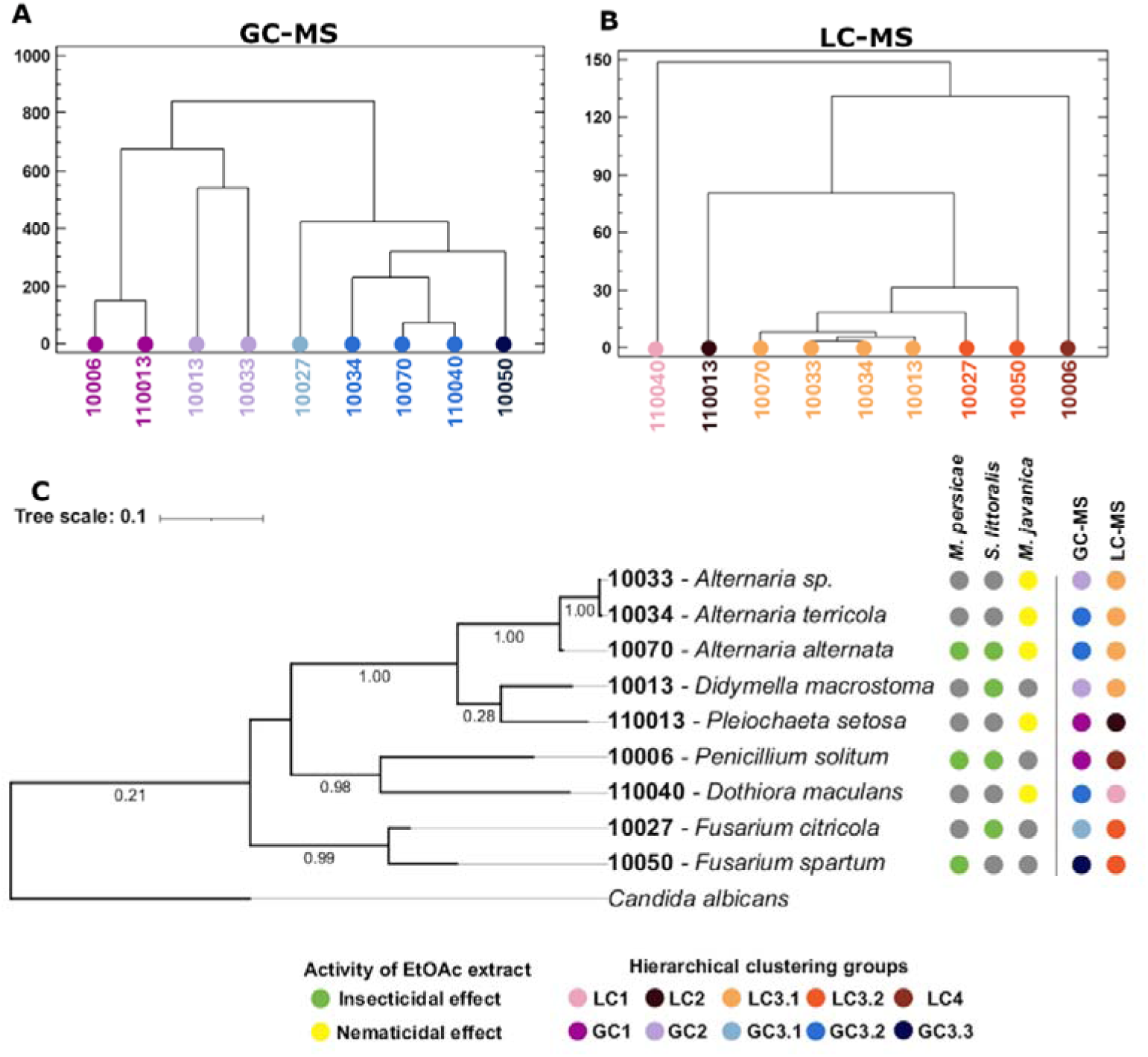
Sample clustering according to metabolomic profiles. Hierarchical clustering of fungal extracts based on their (A) GC-MS (Gas-Chromatography and Mass – Spectrometry) and (B) LC-MS (Liquid Chromatography and Mass – Spectrometry) profile. C) Phylogenetic tree of the fungal isolates based on ITS sequences and classification based on activity of the extracts and the clustering of chromatographic profiles.

A non-target LC-MS analysis showed the presence of at least 28 metabolites, grouped by their retention time, with a range of 7 to 12 metabolites per extract (Supplementary Table S4). Three of the metabolites (No. 130, 134 and 136) were present in all the extracts and two metabolites were absent in just one (No. 132 and 139). The clustering of the results (Figure 2B) showed a profile that could be related to the taxonomical proximity of the isolates. A total of four groups could be established, with three including one species: LC1 (110040, *D. maculans*); LC2 (110013, *P. setosa*); and LC4 (10006, *P. solitum*), and one including three genera, which could be divided into two: LC3.1 (10070-10033-10034, *Alternaria* spp. and 10013, *D. macrostoma*) and LC3.2 (10027-10050, *Fusarium* spp.).

## 4. Discussion

In nature, the preeminent model plant *A. thaliana* hosts a high variety of fungal endophytes^35^. Characterizing its natural endophytic mycobiota offers valuable opportunities to deepen our understanding of plant-endophyte interactions. However, characterization of endophytes isolated from *A. thaliana* is relatively limited, particularly in the context of secondary metabolites biosynthesis and their potential biotechnological applications. In this work, we obtained extracts from 13 *A. thaliana* fungal endophytes and screened them for their activity against the phytoparasitic nematode *M. javanica* and the plant insect pests *M. persicae* and *S. littoralis*. To our knowledge, this is the first study that explores the potential nematicidal and antifeedant effects of secondary metabolites derived from fungal endophytes of *A. thaliana*.

Our findings show that 9 out of the 13 (69%) extracts exhibited nematicidal and/or antifeedant activity, what highlights the enormous potential of fungal endophytes of this model plant to produce bioactive compounds of agricultural interest. The nine isolates with bioactive extracts have different origins regarding population (Polán or Menasalbas) and plant tissue (root, rosette and floral stalk), so their activity is independent from the geographical site or plant organ of provenance. Notably, there where isolates from the roots with activity against leaf pests (i.e. 10006 or 10050) and isolates from the rosette with activity against root nematodes (i.e.10033, 10034 or 110040). Even, the extract from isolate 10070 from the floral stalk was active to both leaf and root attackers. Therefore, the tissue of origin of the isolate did not determine the possible activity of a fungal endophyte.

### 4.1. Nematicidal activity

Isolates 110040 and 110013 from the species *D. maculans* and *P. setosa*, respectively, and the three isolates of *Altern*aria (10033-10034-10070) produced extracts with nematicidal activity. *Dothiora* species are found on plants in terrestrial habitats as saprobes and weak pathogens on stressed plants^48–50^. Previous studies reported the production of hormonemates, compounds with cytotoxic activity against tumoral cells, by endophytic *Dothiora* sp. isolated from *Launaea arborescens*^51^. *P. setosa* is an important pathogenic necrotrophic fungi of grain legumes ^52^. The only compound described for this species is setosol, which has been shown to inhibit the growth of different fungal phytopathogens, such as the fungus *Magnaporthe oryzae* ^53^. However, this is the first report on the nematicidal effects of an organic extract from endophytic species belonging to these two genera.

The fungal genus *Alternaria* is a diverse group of ascomycete fungi which occupy different ecological niches ranging from saprobes to endophytes and pathogens. It is widespread in nature, commonly found in a wide range of hosts and substrates, such as soil, plants, organic matter, wood or textiles^54,55^. From a human health perspective, *Alternaria* is notable due to its airborne spores, which are among the most prevalent allergens^56^. As plant pathogens, *Alternaria* species can cause significant pre- and post-harvest diseases, sometimes leading to mycotoxin accumulation and serious economic losses^57^. Nonetheless, several *Alternaria* species have gained attention for their ability to produce secondary metabolites with a variety of bioactive properties, positioning them as valuable candidates for biotechnological applications in the pharmaceutical and agricultural industries^58,59^. In this work, the three *Alternaria* isolates showed nematicidal activity. Lou et al.^60^ already showed the potential nematicidal effect of endophytic *Alternaria* sp. isolated from *Salvia miltorrhiza*. The bioactive compound was alternariol methyl ether, which was toxic to the model organism *Caenorhabditis elegans* and the plant parasitic nematode *Bursaphelenchus xylophilus*. However, although there is promising evidence of the nematicidal effects of secondary metabolites produced by *Alternaria*, this field remains still underexplored.

### 4.2. Antifeedant activity

Isolate 10006 identified as *P. solitum,* produced an EtOAc extract with antifeedant activity against *M. persicae* and *S. littoralis*. *P. solitum* is known for its role in the spoilage of pome fruits during storage^61^. In addition, this fungus has been isolated from extremophilic environments, such as the acidic waters of the Berkeley Pit Lake^62^ and Antarctica^63^. There is no direct evidence in the literature supporting the use of *P. solitum* metabolites as biopesticides. However, various secondary metabolites from this fungus have exhibited potential bioactive properties. One example is compactin (also referred to as mevastatin), a precursor for the cholesterol-lowering drug pravastatin^64,65^. Additionally, viridicatol, a quinoline alkaloid, exhibited moderate anti-tumor activities against certain cancer cell lines and potent anti-food allergic effects *in vitro*^66^. Nonetheless, many studies have explored the metabolites of other *Penicillium* species for biopesticide purposes. Non endophytic species of this genus are known producers of insecticidal compounds such as tryptoquialanines^67^, indole diketopiperazine alkaloids^68^, okaramine indole alkaloids^69^, meroterpenoids^70^, (-)-botryodiplodin^71^, yaequinolones^72^ or the terpenoid-pyridine oxalicines, active against *S. frugiperda*^73^. The closely related species *P. crustosum,* produces neurotoxic penitrems, which have exhibited insecticidal activity^74^. Furthermore, an endophytic *Penicillium* isolated from *Derris elliptica* also was shown to produce the antifeedant compound rotenone against the lepidopteran *Plutella xylostella* and the aphid *Lipaphis erysimi* ^75^. Still, this is the first report documenting the insecticidal potential of secondary metabolites from *P. solitum* in agriculture.

Isolate 10013, *D. macrostoma,* exhibited antifeedant effects against *S. littoralis*. *D. macrostoma* (formerly *Phoma macrostoma*) is a fungal species with diverse ecological roles. It has been recognized as a plant pathogen, notably causing calyx-end rot in pears during cold storage^76^. However, it has recently shown potential as a biocontrol agent against rapeseed clubroot, significantly reducing disease severity and improving crop yields^77^. The genus *Didymella* is well known for producing a variety of secondary metabolites with phytotoxic and cytotoxic activities, which have garnered attention for their potential as bioherbicides. For instance, an endophytic *Didymella* isolated from mangroves synthesizes cytotoxic ascomylactams (macrocyclic alkaloids), didymetone^78^ and phomapyrrolidones^79^. Additionally, cytotoxic naphthalenones and didymelol have been isolated from the endophytic fungus *D. glomerata*, found in *Saussurea laniceps*^80^. Another example is *D. pinodes*, an aggressive isolate from pea (*Pisum sativum*), which produces pinolidoxin, a phytotoxin affecting several plant species^81^. Despite these findings, this study marks the first report of insect antifeedant effects from an organic extract of endophytic *Didymella*. The compound duroquinone, identified in the GC-MS profile of the EtoAc extract of isolate 10013, may contribute to this effect. Duroquinone has previously been reported to inhibit survival, growth, and pupation in the Black Cutworm (*Agrotis ipsilon*) by reducing ingestion and the efficiency of food conversion^82^. However, further studies are required to confirm its role in the antifeedant activity observed against *S. littoralis*.

Two species of *Fusarium* (isolates 10027-10050) produced active extracts against *M. persicae*. The fungal genus *Fusarium* is a large and significant group of filamentous fungi, primarily found in soil and associated with plants. While certain *Fusarium* species are notorious for causing plant diseases, such as *Fusarium* wilt and root rot in crops like cereals^83–85^, legumes^86^, and bananas^87,88^, as well as for producing harmful mycotoxins^89^, the majority of strains are saprotrophs or endophytes. In fact, *Fusarium* is one of the most abundant endophytic fungal genera, with certain species providing benefits to the plant^90–92^. Moreover, *Fusarium* is a rich source of bioactive compounds from various chemical classes, including those with insecticidal properties^93,94^. Specifically, *F. sambucinum* from *Nicotiana tabacum* produced the prenylated indole alkaloids sclerotiamides and notoamide with potent insecticidal activity against *Helicoverpa*^95^. Moreover, several species of *Fusarium* have been discovered as entomopathogenic in aphids^96^, which could be related with the antifeedant effect observed in their organic extracts.

Apart from the nematicidal effect, EtOAc extract from isolate 10070, identified as *A. alternata,* also showed antifeedant activity against *M. persicae* and *S. littoralis*. Previous studies have reported that *A. alternata* strains isolated from *Azadirachta indica* gave extracts with antifeedant and toxic effects against *S. litura* ^97^. Also, altenuene, an acetyl cholinesterase inhibitor isolated from an endophytic *A. alternata* strain of *Catharantus roseus*, exhibited insecticidal effect against *S. litura*^98^. The EtOAc extract from the 10070 isolate was the only one showing activity against all three plant pests examined in this study, indicating its broad-spectrum efficacy and underscoring its potential as a versatile source of biopesticide products. Additional research will be necessary to identify the specific molecules responsible for this effect.

### 4.3. Analysis of chromatographic profiles

The extracts, analyzed by GC-MS and LC-MS, exhibited a highly diverse composition, with most metabolites being unknown and unique to each extract. Our results did not show a direct correlation between the secondary metabolite profiles and nematicidal or antifeedant activity. Nonetheless, we could observe that two compounds— (No. 34) 2,10-bisaboladien-1-one and (No. 69) tributyl acetylcitrate—were present in all three extracts that demonstrated activity against *M. persicae* (Supplementary Table S3). While acetyltributylcitrate is commonly used as a plasticizer^99^, recent studies have reported it as a naturally occurring component in antimicrobial extracts^100,101^. For example, it has been identified as a bioactive compound in crude extracts of actinomycetes with antibacterial and antifungal properties^101^, as well as in the fungicidal compounds from *Michelia champaca* bark extract^101^. However, its use as a biopesticide remains unexplored. The compound 2,10-bisaboladien-1-one is already known for its antifeedant effects on *M. persicae*, significantly reducing both probing activity and the number of intracellular penetrations^102^. This suggests that 2,10-bisaboladien-1-one may be one of the key compounds responsible for the observed bioactivity.

In addition, no clear relationship between the chemical composition of the extracts and the taxonomical proximity of the isolates was observed. The analysis of the less polar fraction (DCM-soluble) of the extracts, containing the lipid fraction, did not show a taxonomy-dependent distribution. On the other hand, the secondary metabolites in the EtOAc-soluble fraction (analyzed by LC-MS) revealed chemical profiles that could be related to taxonomy, though only to a limited extent. This lack of a clear pattern could be expected, as secondary metabolite profiles in fungi are complex traits influenced by mono- and polyphyletic factors. Although certain metabolites in fungal groups like the *Xylariaceae* family strongly correlate with phylogenies^103^, the inconsistent distribution of secondary metabolites across the fungal kingdom and the great influence of environmental factors in their biosynthesis makes difficult its correlation with phylogeny^104^. Comparative genomic studies have shown that homologous genes and gene clusters related to secondary metabolism are distributed across a wide phylogenetic range of species^105^. In some cases, this distribution is consistent with inheritance from a common ancestor, as seen with the ergot alkaloid gene cluster in *Claviceps* and *Metarhizium*^106^. In other cases, the presence of large, complex gene clusters in distantly related taxa is thought to result from horizontal gene transfer^107^. Within-species genomic comparisons have revealed polymorphisms in secondary metabolite gene clusters, such as gene gain/loss, cluster mobility, and allelic variations, which might explain this evolutionary divergencies^106,108,109^.

In summary, our study underscores the significant potential of fungal endophytes as sources of bioactive compounds with biocontrol capabilities against agricultural pests. The diverse composition of extracts from different fungal isolates does not correlate with the nematicidal and antifeedant activities observed, suggesting similar activities for different metabolites. Nevertheless, identifying the bioactive compounds is essential for gaining a deeper understanding of the mechanisms underlying the observed effects. Future research will focus on bioguided fractionation of selected extracts for the identification of the active compounds. Our results also show that a model host as *A. thaliana*, routinary found in many different anthropic ecosystems in temperate regions^34,35,110^, hosts fungal endophytes with the ability to produce high diversity of compounds that are bioactive. The use of a model plant gives opportunities to better study the relationship of the plant with these endophytes and the conditions in which the secondary metabolites of interest are produced in the interaction with the plants.

## Authors’ contributions

S.D.-G., S.S., A.G.-C. and M.A. conceptualized and designed the research. S.S. provided the fungal isolates. C.G.-S refreshed the isolates and conducted the ITS identification. S.D.-G. constructed the phylogenetic tree, obtained the fungal extracts, conducted the bioassays and prepared the first draft of the manuscript. S.D.-G., A.G.-C. and M.A. prepared the tables and figures. S.D.-G., S.S., A.G.-C. and M.A. contributed to the text of the main manuscript.

All authors have read and agreed to the published version of the manuscript.

## Data availability

All data generated or analysed during this study are included in this published article and its supplementary information files.

## Additional information Competing interests

The authors declare no competing interests

Supplementary information: “Supplemetary Tables.xlsx”

Statement of Exemption from Permission Requirements for the Collection of *Arabidopsis thaliana* Samples: “Statement of Exemption from Permission Requirements.pdf”

## References

1. Chandler, D. et al. The development, regulation and use of biopesticides for integrated pest management. Philosophical Transactions of the Royal Society B: Biological Sciences 366, 1987 (2011).

2. Kumar, J., Ramlal, A., Mallick, D. & Mishra, V. An Overview of Some Biopesticides and Their Importance in Plant Protection for Commercial Acceptance. Plants 2021, Vol. 10, Page 1185 10, 1185 (2021).

3. Macías-Rubalcava, M. L. & Sánchez-Fernández, R. E. Secondary metabolites of endophytic Xylaria species with potential applications in medicine and agriculture. World J Microbiol Biotechnol 33, (2017).

4. Hashem, A. H. et al. Bioactive compounds and biomedical applications of endophytic fungi: a recent review. Microb Cell Fact 22, 1–23 (2023).

5. Hardoim, P. R. et al. The Hidden World within Plants: Ecological and Evolutionary Considerations for Defining Functioning of Microbial Endophytes. Microbiology and Molecular Biology Reviews 79, 293–320 (2015).

6. Lugtenberg, B. J. J., Caradus, J. R. & Johnson, L. J. Fungal endophytes for sustainable crop production. FEMS Microbiol Ecol 92, fiw194 (2016).

7. Yan, L. et al. Beneficial effects of endophytic fungi colonization on plants. Applied Microbiology and Biotechnology vol. 103 3327–3340 Preprint at 10.1007/s00253-019-09713-2 (2019).

8. Mousa, W. K. & Raizada, M. N. The diversity of anti-microbial secondary metabolites produced by fungal endophytes: an interdisciplinary perspective. Front Microbiol 4, 44840 (2013).

9. Ratnaweera, P. B. & de Silva, E. D. Endophytic Fungi: A Remarkable Source of Biologically Active Secondary Metabolites, in Endophytes: Crop Productivity and Protection (eds. Maheshwari, D., Annapurna, K.) 191–212 (Springer, 2017).

10. Bamisile, B. S., Dash, C. K., Akutse, K. S., Keppanan, R. & Wang, L. Fungal endophytes: Beyond herbivore management. Frontiers in Microbiology 9, 544 (2018).

11. Agrios, G. Plant Pathology: Fifth Edition. (Elsevier, 2005).

12. Kusari, S., Hertweck, C. & Spiteller, M. Chemical ecology of endophytic fungi: Origins of secondary metabolites. Chem Biol 19, 792–798 (2012).

13. Schardl, C. et al. Currencies of Mutualisms: Sources of Alkaloid Genes in Vertically Transmitted Epichloae. Toxins (Basel) 5, 1064–1088 (2013).

14. Segaran, G. & Sathiavelu, M. Fungal endophytes: A potent biocontrol agent and a bioactive metabolites reservoir. Biocatal Agric Biotechnol 21, 101284 (2019).

15. Castagnone-Sereno, P., Danchin, E. G. J., Perfus-Barbeoch, L. & Abad, P. Diversity and evolution of root-knot nematodes, genus Meloidogyne: New insights from the genomic era. Annu Rev Phytopathol 51, 203–220 (2013).

16. Abad, P. et al. Genome sequence of the metazoan plant-parasitic nematode Meloidogyne incognita. Nature Biotechnology 26, 909–915 (2008).

17. Azlay, L., El Boukhari, M. E. M., Mayad, E. H. & Barakate, M. Biological management of root-knot nematodes (Meloidogyne spp.): a review. Organic Agriculture 13, 99–117 (2022).

18. Chen, J. xiang & Song, B. an. Natural nematicidal active compounds: Recent research progress and outlook. J Integr Agric 20, 2015–2031 (2021).

19. Van Emden and Harrington. Aphids as Crop Pests. (CABI, UK, 2017).

20. Ali, J. et al. Peach–Potato Aphid Myzus persicae: Current Management Strategies, Challenges, and Proposed Solutions. Sustainability 15, 11150 (2023).

21. Pasiecznik, N. M. et al. CABI/EPPO distribution maps of plant pests and plant diseases and their important role in plant quarantine. EPPO Bulletin 35, 1–7 (2005).

22. Cheng, T. et al. Genomic adaptation to polyphagy and insecticides in a major East Asian noctuid pest. Nature Ecology & Evolution 1, 1747–1756 (2017).

23. Hilliou, F., Chertemps, T., Maïbèche, M. & Le Goff, G. Resistance in the genus Spodoptera: Key insect detoxification genes. Insects 12, 544 (2021).

24. Andrés, M. F., Diaz, C. E., Giménez, C., Cabrera, R. & González-Coloma, A. Endophytic fungi as novel sources of biopesticides: the Macaronesian Laurel forest, a case study. Phytochemistry Reviews 16, 1009–1022 (2017).

25. Bogner, C. W. et al. Bioactive secondary metabolites with multiple activities from a fungal endophyte. Microb Biotechnol 10, 175–188 (2017).

26. Kaushik, N. et al. Chemical Composition of an Aphid Antifeedant Extract from an Endophytic Fungus, Trichoderma sp. EFI671. Microorganisms 8, 420 (2020).

27. Diaz, C. E. et al. Antifeedant, antifungal and nematicidal compounds from the endophyte Stemphylium solani isolated from wormwood. Scientific Reports 14, 1–10 (2024).

28. Reyes Castillo, N., et al. Optimization of fungicidal and acaricidal metabolite production by endophytic fungus Aspergillus sp. SPH2. Bioresour Bioprocess 11, 1–12 (2024).

29. Poveda, J., Díaz-González, S., Díaz-Urbano, M., Velasco, P. & Sacristán, S. Fungal endophytes of Brassicaceae: Molecular interactions and crop benefits. Front Plant Sci 13, 932288 (2022).

30. Koornneef, M. & Meinke, D. The development of Arabidopsis as a model plant. The Plant Journal 61, 909–921 (2010).

31. Hiruma, K. et al. Root Endophyte Colletotrichum tofieldiae Confers Plant Fitness Benefits that Are Phosphate Status Dependent. Cell 165, 464–474 (2016).

32. Díaz-González, S. et al. Mutualistic Fungal Endophyte Colletotrichum tofieldiae Ct0861 Colonizes and Increases Growth and Yield of Maize and Tomato Plants. Agronomy 10, 1493 (2020).

33. Díaz-González, S. et al. Plant Growth Promoting fungal endophyte Colletotrichum tofieldiae Ct0861 reduces mycotoxigenic Aspergillus fungi in maize grains. bioRxiv 2025.01.17.633531 (2025) doi:10.1101/2025.01.17.633531.

34. Mesny, F. et al. Genetic determinants of endophytism in the Arabidopsis root mycobiome. Nature Communications 12:1 12, 1–15 (2021).

35. García, E., Alonso, Á., Platas, G. & Sacristán, S. The endophytic mycobiota of Arabidopsis thaliana. Fungal Divers 60, 71–89 (2013).

36. Bokulich, N. A. & Mills, D. A. Improved selection of internal transcribed spacer-specific primers enables quantitative, ultra-high-throughput profiling of fungal communities. Appl Environ Microbiol 79, 2519–2526 (2013).

37. Doyle, J. J. & Doyle, J. L. A rapid DNA isolation procedure for small quantities of fresh leaf tissue. PHYTOCHEMICAL BULLETIN (1987).

38. Katoh, K. & Standley, D. M. MAFFT Multiple Sequence Alignment Software Version 7: Improvements in Performance and Usability. Mol Biol Evol 30, 772–780 (2013).

39. Capella-Gutiérrez, S., Silla-Martínez, J. M. & Gabaldón, T. trimAl: a tool for automated alignment trimming in large-scale phylogenetic analyses. Bioinformatics 25, 1972–1973 (2009).

40. Price, M. N., Dehal, P. S. & Arkin, A. P. FastTree 2 - Approximately maximum-likelihood trees for large alignments. PLoS One 5, e9490 (2010).

41. Letunic, I. & Bork, P. Interactive Tree of Life (iTOL) v6: recent updates to the phylogenetic tree display and annotation tool. Nucleic Acids Res 52, W78–W82 (2024).

42. Moo-Koh, F. A. et al. In Vitro Assessment of Organic and Residual Fractions of Nematicidal Culture Filtrates from Thirteen Tropical Trichoderma Strains and Metabolic Profiles of Most-Active. Journal of Fungi 8, 82 (2022).

43. Schneider-Orelli, O. Entomologisches Praktikum: Einführung in Die Land-Und. Forstwirtschaftliche Insektenkunde. (1947).

44. Andrés, M. F., González-Coloma, A., Sanz, J., Burillo, J. & Sainz, P. Nematicidal activity of essential oils: A review. Phytochemistry Reviews 11, 371–390 (2012).

45. Poitout, S. & Bues, R. Rearing of several species of Lepidoptera Noctuidae on a rich artificial medium and a simplified artificial medium. Annales de Zoologie Ecologie Anim ale 2, 79–91 (1970).

46. Gonzalez-Coloma, A. et al. Structure- and Species-Dependent Insecticidal Effects of neo-Clerodane Diterpenes. J Agric Food Chem 48, 3677–3681 (2000).

47. Schneider, C. A., Rasband, W. S. & Eliceiri, K. W. NIH Image to ImageJ: 25 years of image analysis. Nature Methods 9, 671–675 (2012).

48. Crous, P. W. & Groenewald, J. Z. They seldom occur alone. Fungal Biol 120, 1392–1415 (2016).

49. Crous, P. W. & Groenewald, J. Z. The Genera of Fungi - G 4: Camarosporium and Dothiora. IMA Fungus 8, 131–152 (2017).

50. Hyde, K. D. et al. The Numbers of Fungi: Is the Descriptive Curve Flattening? Fungal Diversity 103, 219–271 (Springer Netherlands, 2020).

51. Pérez-Bonilla, M. et al. Hormonemate Derivatives from Dothiora sp., an Endophytic Fungus. J Nat Prod 80, 845–853 (2017).

52. Agudo-Jurado, F. J., Reveglia, P., Rubiales, D., Evidente, A. & Barilli, E. Status of Phytotoxins Isolated from Necrotrophic Fungi Causing Diseases on Grain Legumes. International Journal of Molecular Sciences 24, 5116 (2023).

53. Okeke, B., Seigle-Murandi, F., Steiman, R. & Kaouadji, M. Setosol, a Biologically Active Heptaketide-like Metabolite from the Pleiochaeta setosa Phytopathogen. Biosci Biotechnol Biochem 58, 734–736 (1994).

54. Thomma, B. P. H. J. Alternaria spp.: from general saprophyte to specific parasite. Mol Plant Pathol 4, 225–236 (2003).

55. Lawrence, D. P., Rotondo, F. & Gannibal, P. B. Biodiversity and taxonomy of the pleomorphic genus Alternaria. Mycological Progress 15, 1–22 (2015).

56. Kasprzyk, I. et al. Air pollution by allergenic spores of the genus Alternaria in the air of central and eastern Europe. Environmental Science and Pollution Research 22, 9260–9274 (2015).

57. Pinto, V. E. F. & Patriarca, A. Alternaria Species and Their Associated Mycotoxins. Methods in Molecular Biology 1542, 13–32 (2017).

58. Zhao, S. et al. Secondary metabolites of Alternaria: A comprehensive review of chemical diversity and pharmacological properties. Front Microbiol 13, 1085666 (2023).

59. Lou, J., Fu, L., Peng, Y. & Zhou, L. Metabolites from Alternaria Fungi and Their Bioactivities. Molecules 18, 5891–5935 (2013).

60. Lou, J. et al. Alternariol 9-methyl ether from the endophytic fungus Alternaria sp. Samif01 and its bioactivities. Brazilian Journal of Microbiology 47, 96–101 (2016).

61. Jurick, W. M. et al. Penicillium solitum produces a polygalacturonase isozyme in decayed Anjou pear fruit capable of macerating host tissue in vitro. Mycologia 104, 604–612 (2012).

62. Stierle, D. B., Stierle, A. A., Girtsman, T., McIntyre, K. & Nichols, J. Caspase-1 and -3 inhibiting drimane sesquiterpenoids from the extremophilic fungus Penicillium solitum. J Nat Prod 75, 262–266 (2012).

63. Gonçalves, V. N. et al. Penicillium solitum: A mesophilic, psychrotolerant fungus present in marine sediments from Antarctica. Polar Biol 36, 1823–1831 (2013).

64. Boruta, T., Przerywacz, P., Ryngajllo, M. & Bizukojc, M. Bioprocess-related, morphological and bioinformatic perspectives on the biosynthesis of secondary metabolites produced by Penicillium solitum. Process Biochemistry 68, 12–21 (2018).

65. Frisvad, J. C. & Filtenborg, O. Terverticillate Penicillia: Chemotaxonomy and Mycotoxin Production. Mycologia 81, 837–861 (1989).

66. He, Z. H. et al. Chemical constituents of the deep-sea-derived penicillium solitum. Mar Drugs 19, 580 (2021).

67. Costa, J. H. et al. Monitoring indole alkaloid production by Penicillium digitatum during infection process in citrus by Mass Spectrometry Imaging and molecular networking. Fungal Biol 123, 594–600 (2019).

68. Jia, B., Ma, Y., Chen, D., Chen, P. & Hu, Y. Studies on Structure and Biological Activity of Indole Diketopiperazine Alkaloids. Progress in Chemistry 30, 1067–1081 (2018).

69. Hayashi, H., Takiuchi, K., Murao, S. & Arai, M. Structure and insecticidal activity of new indole alkaloids, okaramines a and b, from penicillium simplicissimum ak-40. Agric Biol Chem 53, 461–469 (1989).

70. Geris, R., Rodrigues-Fo, E., du Silva, H. H. G. & da Silva, I. G. Larvicidal Effects of Fungal Meroterpenoids in the Control of Aedes aegypti L., the Main Vector of Dengue and Yellow Fever. Chem Biodivers 5, 341–345 (2008).

71. Cabedo, N., López-Gresa, M. P., Primo, J., Ciavatta, M. L. & González-Mas, M. C. Isolation and structural elucidation of eight new related analogues of the mycotoxin (-)- botryodiplodin from Penicillium coalescens. J Agric Food Chem 55, 6977–6983 (2007).

72. Uchida, R., Imasato, R., Tomoda, H. & Omura, S. Yaequinolones, New Insecticidal Antibiotics Produced by Penicillium sp. FKI-2140. The Journal of Antibiotics 59, 652–658 (2006).

73. Li, C., Gloer, J. B., Wicklow, D. T. & Dowd, P. F. Antiinsectan decaturin and oxalicine analogues from Penicillium thiersii. J Nat Prod 68, 319–322 (2005).

74. Carmen González, M., et al. Insecticidal Activity of Penitrems, Including Penitrem G, a New Member of the Family Isolated from Penicillium crustosum. J Agric Food Chem 51, 2156– 2160 (2003).

75. Hu, M. Y. et al. Insecticidal metabolites produced by Penicillium spp., an endophytic fungus in Derris elliptica Benth. Allelopathy Journal 21, 349–360 (2008).

76. Wenneker, M., Pham, K. T. K. & Kots, K. First Report of Didymella macrostoma Causing Calyx-End Rot of Pear (Pyrus communis) in the Netherlands. Plant Disease, 107, 2855 (2023).

77. Li, G. et al. Biological Control of Rapeseed Clubroot (Plasmodiophora brassicae) Using the Endophytic Fungus Didymella macrostoma P2. Plant Disease, 1088, 2399–2409 (2024).

78. Yuan, Y., Wang, G., She, Z., Chen, Y. & Kang, W. Metabolites isolated from the mangrove endophytic fungus Didymella sp. CYSK-4 and their cytotoxic activities. Fitoterapia 171, 105692 (2023).

79. Chen, Y., et al. Ascomylactams A-C, Cytotoxic 12- or 13-Membered-Ring Macrocyclic Alkaloids Isolated from the Mangrove Endophytic Fungus Didymella sp. CYSK-4, and Structure Revisions of Phomapyrrolidones A and C. J Nat Prod 82, 1752–1758 (2019).

80. Luo, G. et al. Naphthalenones and Naphthols Isolated from the Saussurea laniceps Endophytic Fungus Didymella glomerata X223. Chem Biodivers 17, e2000315 (2020).

81. Cimmino, A. et al. Pinolide, a new nonenolide produced by Didymella pinodes, the causal agent of ascochyta blight on Pisum sativum. J Agric Food Chem 60, 5273–5278 (2012).

82. Reese, J. C. & Beck, S. D. Effects of Allelochemics on the Black Cutworm, Agrotis ipsilon; Effects of p-Benzoquinone, Hydroquinone, and Duroquinone on Larval Growth, Development, and Utilization of Food. Ann Entomol Soc Am 69, 59–67 (1976).

83. Yli-Mattila, T. Ecology and evolution of toxigenic Fusarium species in cereals in northern Europe and Asia. Journal of Plant Pathology 92, 7–18 (2010).

84. Arata, A. F. et al. The richness of Fusarium species in maize tassels and their relationship with Fusarium stalk rot. Eur J Plant Pathol 168, 351–362 (2024).

85. Zuo, S. et al. Assessment of Genetic Diversity and the Population Structure of Species from the Fusarium fujikuroi Species Complex Causing Fusarium Stalk Rot of Maize. Journal of Fungi 10, 574 (2024).

86. Sampaio, A. M., De Sousa Araújo, S., Rubiales, D. & Patto, M. C. V. Fusarium Wilt Management in Legume Crops. Agronomy 10, 1073 (2020).

87. Dita, M., Barquero, M., Heck, D., Mizubuti, E. S. G. & Staver, C. P. Fusarium wilt of banana: Current knowledge on epidemiology and research needs toward sustainable disease management. Front Plant Sci 871, 398832 (2018).

88. Ploetz, R. C. Fusarium wilt of banana. Phytopathology 105, 1512–1521 (2015).

89. Ji, F. et al. Occurrence, toxicity, production and detection of Fusarium mycotoxin: a review. Food Production, Processing and Nutrition 1, 1–14 (2019).

90. Pappas, M. L. et al. The beneficial endophytic fungus fusarium solani strain K alters tomato responses against spider mites to the benefit of the plant. Front Plant Sci 9, 408405 (2018).

91. Vu, T., Hauschild, R. & Sikora, R. A. Fusarium oxysporum endophytes induced systemic resistance against Radopholus similis on banana. Nematology 8, 847–852 (2006).

92. de Lamo, F. J. & Takken, F. L. W. Biocontrol by Fusarium oxysporum Using Endophyte-Mediated Resistance. Front Plant Sci 11, 500488 (2020).

93. Ahmed, A. M., Mahmoud, B. K., Millán-Aguiñaga, N., Abdelmohsen, U. R. & Fouad, M. A. The endophytic Fusarium strains: a treasure trove of natural products. RSC Adv 13, 1339– 1369 (2023).

94. Toghueo, R. M. K. Bioprospecting endophytic fungi from Fusarium genus as sources of bioactive metabolites. Mycology 11, 1–21 (2020).

95. Zhang, P. et al. Angularly Prenylated Indole Alkaloids with Antimicrobial and Insecticidal Activities from an Endophytic Fungus Fusarium sambucinum TE-6L. J Agric Food Chem 67, 11994–12001 (2019).

96. Santos, A. C. da S., Diniz, A. G., Tiago, P. V. & Oliveira, N. T. de. Entomopathogenic Fusarium species: a review of their potential for the biological control of insects, implications and prospects. Fungal Biol Rev 34, 41–57 (2020).

97. Kaur, T., Kaur, J., Kaur, A. & Kaur, S. Larvicidal and growth inhibitory effects of endophytic Aspergillus niger on a polyphagous pest, Spodoptera litura. Phytoparasitica 44, 465–476 (2016).

98. Bhagat, J. et al. Cholinesterase inhibitor (Altenuene) from an endophytic fungus Alternaria alternata: optimization, purification and characterization. J Appl Microbiol 121, 1015–1025 (2016).

99. Chaos, A. et al. Tributyl citrate as an effective plasticizer for biodegradable polymers: effect of plasticizer on free volume and transport and mechanical properties. Polym Int 68, 125– 133 (2019).

100. Awad, N. M., Rasmey, A.-H. M., Elshamy, A. I. & Aboseidah, A. Antimicrobial Activities of Some Actinomycetes Isolated from Cultivated Soil, Egypt. Frontiers in Scientific Research and Technology 8, (2024).

101. Bawa, I. G. A. G., Santi, S. R., Rita, W. S., Suryanadi, O. & Indyan, G. Active compounds of Michelia champaca bark extract against Curvularia verruculosa fungi causing leaf spot disease in rice (Oryza sativa L.). Journal of Applied and Natural Science 16, 420–426 (2024).

102. Gutiérrez, C., Fereres, A., Reina, M., Cabrera, R. & González-Coloma, A. Behavioral and sublethal effects of structurally related lower terrenes on Myzus persicae. J Chem Ecol 23, 1641–1650 (1997).

103. Stadler, M. Importance of secondary metabolites in the Xylariaceae as parameters for. Current Research in Environmental & Applied Mycology 1, 75–133 (2011).

104. Frisvad, J. C., Andersen, B. & Thrane, U. The use of secondary metabolite profiling in chemotaxonomy of filamentous fungi. Mycol Res 112, 231–240 (2008).

105. Spatafora, J. W. & Bushley, K. E. Phylogenomics and evolution of secondary metabolism in plant-associated fungi. Curr Opin Plant Biol 26, 37–44 (2015).

106. Young, C. A., et al. Genetics, Genomics and Evolution of Ergot Alkaloid Diversity. Toxins 2015, Vol. 7, Pages 1273-1302 7, 1273–1302 (2015).

107. Wisecaver, J. H., Slot, J. C. & Rokas, A. The Evolution of Fungal Metabolic Pathways. PLoS Genet 10, e1004816 (2014).

108. Liu, L., Xi, Z. & Davis, C. C. Coalescent Methods Are Robust to the Simultaneous Effects of Long Branches and Incomplete Lineage Sorting. Mol Biol Evol 32, 791–805 (2015).

109. Lind, A. L. et al. Drivers of genetic diversity in secondary metabolic gene clusters within a fungal species. PLoS Biol 15, e2003583 (2017).

110. Thiergart, T. et al. Root microbiota assembly and adaptive differentiation among European Arabidopsis populations. Nat Ecol Evol 4, 122–131 (2019).

